# Intracellular *Pseudomonas aeruginosa* within the airway epithelium of Cystic Fibrosis lung tissues

**DOI:** 10.1101/2023.08.17.552973

**Authors:** Karim Malet, Emmanuel Faure, Damien Adam, Jannik Donner, Lin Liu, Sarah-Jeanne Pilon, Richard Fraser, Peter Jorth, Dianne K. Newman, Emmanuelle Brochiero, Simon Rousseau, Dao Nguyen

## Abstract

**RATIONALE:** *Pseudomonas aeruginosa* (*P.a*.) is the major bacterial pathogen colonizing the airways of adult cystic fibrosis (CF) patients and causes chronic infections that persist despite antibiotic therapy. Intracellular bacteria may represent an unrecognized reservoir of bacteria that evades the immune system and antibiotic therapy. While the ability of *P.a*. to invade and survive within epithelial cells has been described *in vitro* in different epithelial cell models, evidence of this intracellular lifestyle in human lung tissues is currently lacking.

**OBJECTIVES:** To detect and characterize intracellular *P.a*. in CF airway epithelium from human lung explant tissues.

**METHODS:** We sampled the lung explant tissues from CF and non-CF patients undergoing lung transplantation and analyzed lung tissue sections for the presence of intracellular *P.a*. by quantitative culture and microscopy, in parallel to histopathology and airway morphometry.

**MEASUREMENTS AND MAIN RESULTS:** *P.a*. was isolated from the lungs of 7 CF patients undergoing lung transplantation. Microscopic assessment revealed the presence of intracellular *P.a*. within airway epithelial cells in 3 out of the 7 lungs analyzed, at a varying but rare frequency. We observed those events occurring in lung regions with high bacterial burden.

**CONCLUSION:** This is the first study describing the presence of intracellular *P.a*. in CF lung tissues. While intracellular *P.a*. in airway epithelial cells are likely relatively rare events, our findings highlight the plausible occurrence of this intracellular bacterial reservoir in chronic CF infections.

## Introduction

Due to impaired mucociliary clearance and other defects in host defenses linked to CFTR dysfunction, individuals with the genetic disease cystic fibrosis (CF) are highly susceptible to bacterial colonization and chronic airway infections. By adulthood, the majority of individuals with CF have developed chronic airway infections with the opportunistic Gram-negative bacterial pathogen *Pseudomonas aeruginosa*. Even with antibiotic eradication therapy given at the first evidence of detectable bacteria in sputum cultures, *P*.*a*. can progress to a chronic and persistent infection that resist eradication by host immune defenses and antibiotic therapies (1, 2) and is associated with increased morbidity (3). In fact, even with the development of highly efficient CFTR modulator therapies, CF airways are rarely cleared of *P*.*a*., suggesting that restoration of CFTR-dependent mucociliary clearance and host defenses is not sufficient to eradicate *P*.*a*. (4–7).

Previous microscopy studies of CF lung explant and post-mortem tissues reported that *P*.*a*. was predominantly found as microcolonies within the mucus overlaying the bronchial epithelium and obstructing small airways, and was rarely reported to adhere to airway epithelium (8–12). While these studies support the widely accepted notion that *P*.*a*. is an extracellular pathogen, several groups including ours have reported on the ability of *P*.*a*, to invade and survive intracellularly within epithelial cells using *in vitro* infection models of cornea, lung or urinary tract epithelium. *P*.*a*. hijacks the host cell internalization machinery following interaction of adhesins with receptors on target cell surface (13, 14). The host cytoskeleton, tyrosine kinases (15–18) and phosphoinositides (19, 20) are recruited, allowing bacterial entry. Distinct intracellular niches have been reported, ranging from intravacuolar dwelling (20–22) to cytosolic release and further survival in non-apoptotic blebs (23–25). Bacterial factors such as the type 3 secretion system also influence the fate of intracellular *P*.*a*., highlighting its strain dependent variable capacity to persist intracellularly (21, 25–27).

Intracellular bacteria are protected from phagocytic clearance and antimicrobial therapies that exhibit poor diffusion into mammalian cells, and could thus represent a reservoir that contributes to the persistence of *P*.*a*. infections. While this hypothesis is compelling, *in vivo* evidence of intracellular *P*.*a*. has been extremely limited. Using murine models of infection, intracellular *P*.*a*. has only been reported in keratinocytes for corneal infections, and bladder epithelial cells for urinary tract infections (22, 25, 28, 29).

To date, no studies have thoroughly assessed CF lung tissues for evidence of intracellular *P*.*a*. (12, 30). In this study, we aim to fill this gap by analyzing lung explant tissues from a cohort of CF patients undergoing lung transplantation to detect for the presence of intracellular *P*.*a*. and characterize their distribution within the airway epithelium. Through culture and immunodetection-based methods combined with light and fluorescence microscopy, our analyses reveal that intracellular *P*.*a*,. although relatively rare, can be detected within airway epithelial cells. Moreover, morphometric analyses of airway thin sections and immunofluorescence of optically cleared thick lung tissue sections further reveal the spatial context of intracellular *P*.*a*. within CF airways.

### Material and methods

Detailed methods for tissue sample collection, microbiological analyses, tissue clearing, immunofluorescence and immunohistochemistry staining, histology and confocal imaging are provided in the online supplements.

### Lung tissues and clinical data collection

Human lung tissues were sampled from lung explants obtained from CF patients undergoing lung transplantation and from a healthy lung donor at the Centre Hospitalier de l‘Université de Montréal (CHUM). Fresh lung explants were kept in sterile normal saline at 4°C immediately following surgical resection and sampled within 18 hrs. In each CF patient, one explanted lung was sampled (left or right) by sterile dissection, with at least one specimen (∼1.5 g) per lobe and up to 4 specimens per patient lung. Each tissue specimen was further cut into three contiguous samples (range 140 – 1800 mg) and placed in either sterile phosphate buffer saline (PBS) for microbiological analyses, 4% paraformaldehyde (PFA) for tissue clearing, or 10% formalin for histology and immunohistochemistry (IHC). For the healthy lung donor, airway tissue samples were taken at the bronchial resection site. Data on the CFTR genotype, age, clinical microbiological data, current antibiotic treatment were obtained for each CF patient from the CHUM electronic medical records and the Canadian Cystic Fibrosis Registry. All sputum cultures were performed by the CHUM clinical microbiology lab as part of standard clinical care. All tissues and clinical data were obtained following patient informed consent through the CR-CHUM Respiratory Tissue and Cell Biobank using REB approved protocols #CE-08.063 at the CHUM and #2018-3582 at the McGill University Health Centre.

### Culture-based microbiological analysis of lung tissues

Isolation of extracellular and intracellular bacteria was performed from tissues stored in PBS on LB agar (Difco 244620), Pseudomonas isolation agar (PIA, BD #292710), Blood agar (BA, BD #254071), Mannitol salt agar (MSA, BD 211407) and McConkey agar (MCA, BD 299769) plates.

### Immunohistochemistry (IHC) for P.a. detection and histopathology scoring of lung tissues

IHC staining of *P*.*a*. was performed using a rabbit anti-*Pseudomonas* polyclonal antibody (Abcam, ab68538) and Hematoxylin counterstain. For histopathology scoring, thin sections were stained with Hematoxylin and Eosin. All stained lung thin sections were analyzed in a blinded manner for *P*.*a*. quantification and histology. See online supplements for details.

### Lung tissue clearing

Lung tissues were cleared by passive MiPACT methods as described in (31). Sections were stained with mouse anti-*P. aeruginosa* monoclonal (Thermo Fisher, # MA183430) and then with goat anti-mouse Alexa Fluor 555 (Thermo Fisher, # A32727) antibodies, Alexa-488 conjugated wheat-germ agglutinin (WGA, Thermo Fisher, #W11261) and DAPI. See online supplements for details.

### Statistical Analysis

Statistical analysis was performed using Prism 9 software (GraphPad). Means +/- standard deviations (SD) were calculated and displayed unless stated otherwise. Comparisons were performed using a Mann-Whitney U test, significance was accepted at *P*-value less than 0.05.

### Data availability

The data supporting our results are shown in the main and supplemental figures, as indicated in the manuscript. Raw IHC images used for analyses are available at https://omero.med.ualberta.ca/webclient/?show=dataset-1706

## Results

We first report the case of a CF patient (CF1, delF508 / 711+1G CFTR mutations) who underwent bilateral lung transplantation. Serial sputum cultures performed up to 27 months prior to transplantation as part of standard clinical care revealed colonization by *Stenotrophomonas maltophilia* and *Aspergillus fumigatus*. During the same period, only one out of 15 sputum cultures performed was positive for *P*.*a*. at 6 months prior to transplantation (Figure 1A and Table E1). Immediately after surgical resection, the lung explant was sampled for microbiological analysis and microscopy.

**Figure 1.**
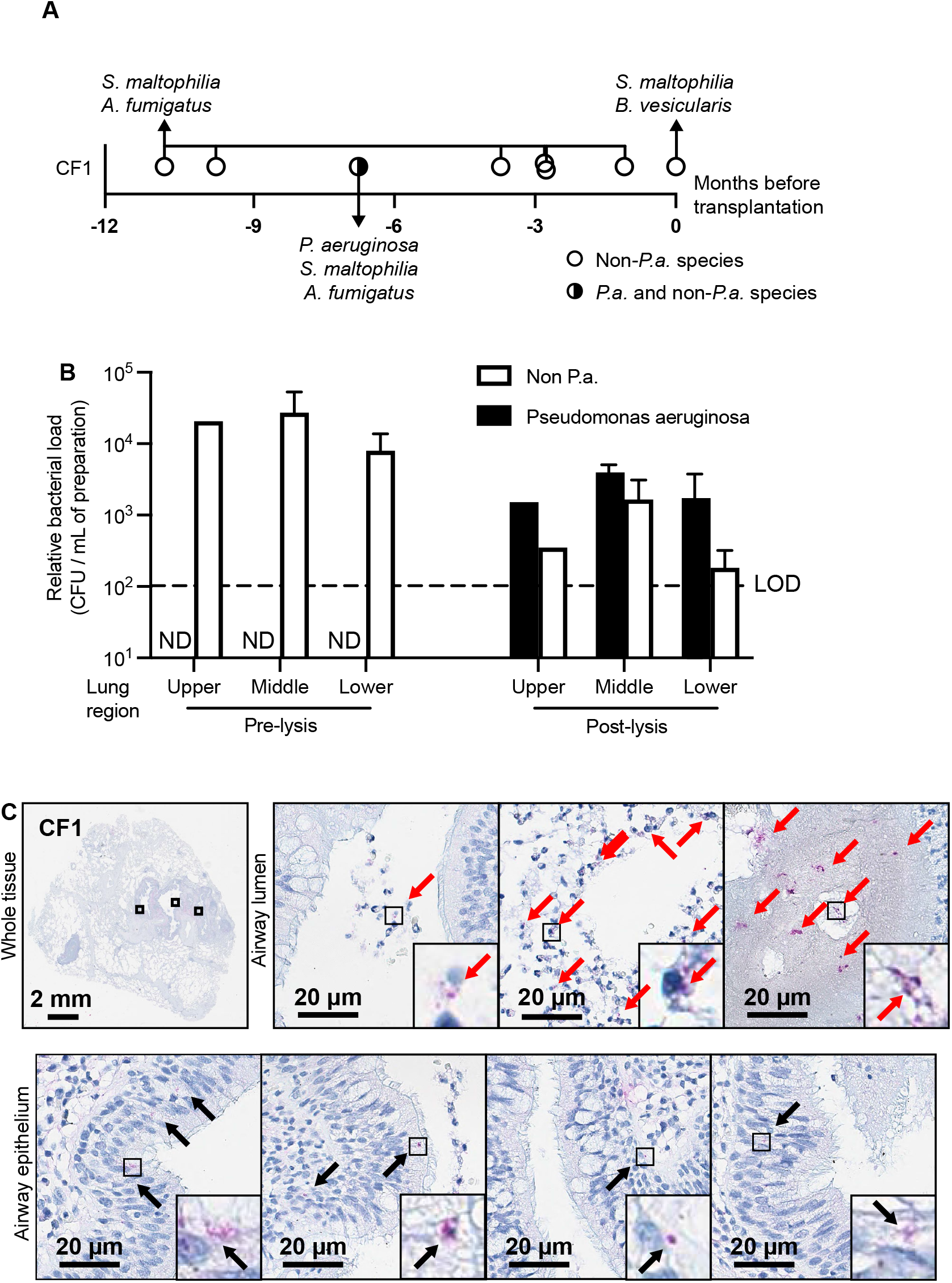
A case study of intracellular *Pseudomonas aeruginosa (P.a.)* in a CF lung explant. (A) Timeline of sputum culture microbiology from patient CF1 collected up to 1 year prior to lung transplantation. Detailed microbiology results are available in Table E1. (B) Microbiological analysis of CF1 lung tissue with cultures performed in pre-lysis and post-lysis samples. Relative viable bacterial load for *P*.*a*. and non-*P*.*a*. species are indicated. Results are shown as mean +/- SD of 2 biological replicates. LOD limit of detection. (C) Representative IHC microscopy images of CF1 explant tissues (whole tissue section, airway lumen and airway epithelium) stained with hematoxylin and anti-*P*.*a*. polyclonal antibody. High magnification images are shown in the insets. IHC bacterial signals are indicated with arrows in red (*P*.*a*. in airway lumen) or black (intracellular *P*.*a*. in airway epithelial cells).

After tissue resuspension in an equal volume of PBS, extracellular bacterial burdens of *P*.*a*. and other bacterial species were assessed by plating PBS washes on selective and non-selective media, respectively. Consistent with the concurrent sputum cultures, only non-*P*.*a*. species were cultured from the three lobes sampled (averages ranging from 8 to 27.3 × 10^4 CFU/mL) (Figure 1B). Tissue samples were then treated with antibiotics that diffuse poorly inside eukaryotic cells (tobramycin and colistin), and lysed to generate ”post-lysis” samples enriched for bacteria protected from the antibiotic cocktail, including intracellular bacteria. As expected, non-*P*.*a*. species were cultured from ”post-lysis” samples, albeit displaying a 16- to 69-fold lower bacterial burden (range 0.09 to 2.7×10^4 CFU/mL) compared to the corresponding pre-lysis samples. Surprisingly, we also retrieved viable *P*.*a*. from the same ”post-lysis” samples (range 0.3 to 3.2×10^4 CFU/mL), while no *P*.*a*. was recovered from the ”pre-lysis” samples (limit of detection 10^2 CFU/mL, Figure 1B). *P*.*a*. clones isolated from CF lungs are described in Table E2. Together, these observations suggested the existence of unrecognized bacterial reservoirs that were not readily sampled in airway secretions (sputum samples) or tissue washes, and thus raised the possibility of an intracellular *P*.*a*. reservoir in these lung explants.

We next explored the possibility of intracellular *P*.*a*. within airway epithelial cells. We performed immunohistochemistry analysis of lung explant sections from patient CF1 using a polyclonal anti-*P*.*a*. antibody. We first established staining controls for lung tissue (Supplemental Figure E1A-F) and validated the antibody’s specificity for *P*.*a*. and its ability to recognize *P*.*a*. CF clinical isolates (Supplemental Figure E2A-C). Consistent with previous studies (9–12, 30, 32), we detected *P*.*a*. microcolony aggregates (size 3-5 μm) within mucopurulent secretions as well as individual bacteria associated to eukaryotic cells in the airway lumen (most likely macrophages or PMN) (Figure 1C). Most interesting, we also observed rare intracellular signals consistent with single bacteria within ciliated airway epithelial cells (Figure 1C). The microbiological and IHC findings in CF1 suggested that intracellular *P*.*a*. could be observed within airway epithelial cells, albeit at very low bacterial burden.

These initial observations in patient CF1 led us to examine the lung explant tissues of 6 additional CF patients for further evidence of intracellular *P*.*a*., and tissue sections from one non-CF patient as control. The clinical characteristics of all CF patients are presented in Table 1. All CF lung explants were processed as described in the workflow for histology and IHC (Figure 2). The results from sputum cultures performed prior to the transplantation as well as bacterial cultures of ”pre-lysis” and ”post-lysis” samples (as done for CF1) are reported in Figure 3A and Table E2. All patients C2 to C7 had evidence of remote *P*.*a*. infection (up to 14 months before transplantation). Although 3 out of 6 patients had *P*.*a*.*-*negative sputum cultures at the time of transplantation, *P*.*a*. was cultured from pre-lysis and post-lysis” explant tissue samples from all CF2 to CF7 patients (Figure 3A).

**Table 1.** Characteristics of the CF patients (n=7) at the time of lung transplantation. *Based on the isolation of any *P*.*a*. (with or without other non-*P*.*a*. species).

| **Characteristic** |  |
| --- | --- |
| Female (%) | 57 |
| Average age (years) | 30 |
| CFTR genotype |  |
| Homozygous delF508 (%) | 57 |
| Heterozygous delF508 (%) | 29 |
| Other (%) | 14 |
| Average lung function FEV<sub>1</sub> (% predicted) | 27.3 |
| *P. aeruginosa*-positive sputum (% of all patients) |  |
| Before transplantation (%) * | 43 |
| 12 months prior to transplantation (%) * | 86 |

**Figure 2.**
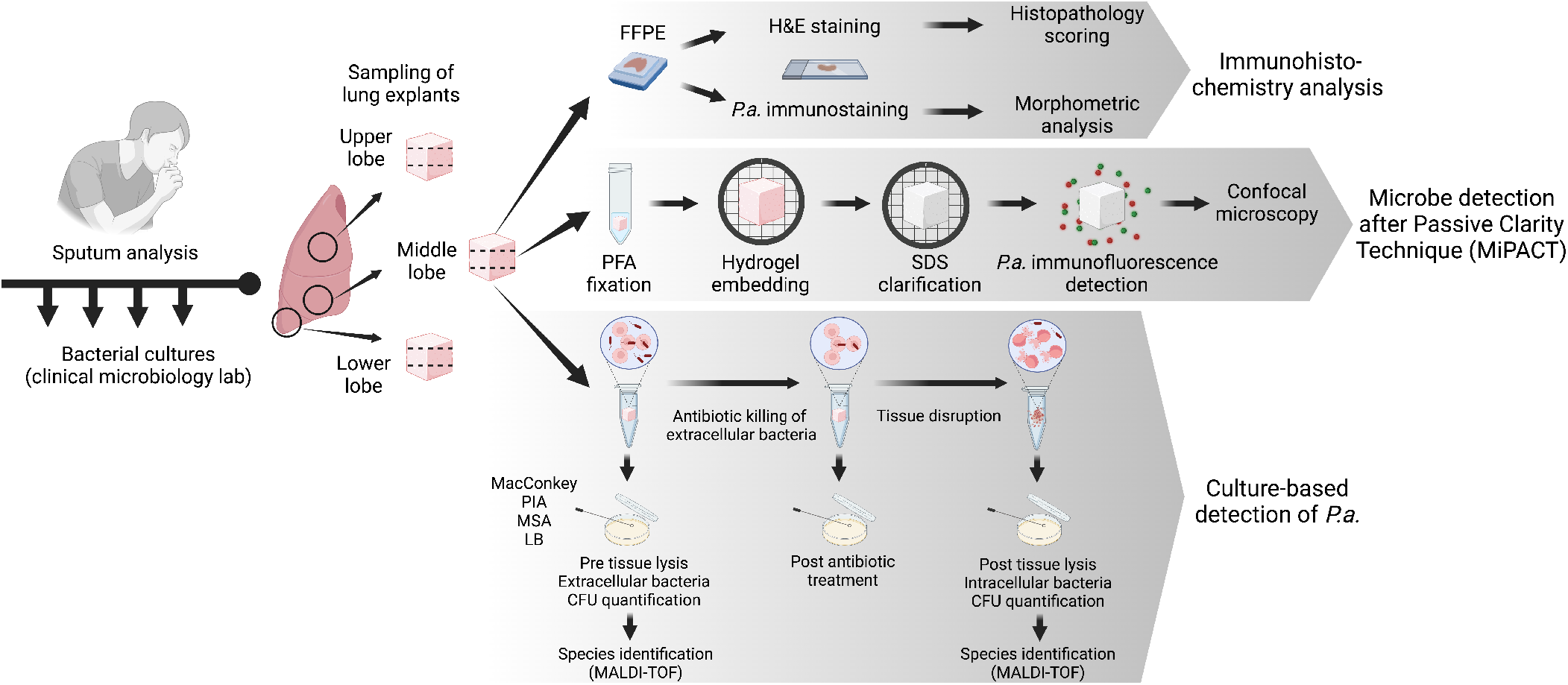
Workflow for sampling, processing and analyses of CF lung explants. Each lung explant were processed for parallel culture-based bacterial detection, IHC and histopathology. A subset of lung explants were analyzed by MiPACT (Microbial detection after Passive Clarity technique) coupled to immunofluorescence. FFPE formalin fixed paraffin embedded.

**Figure 3.**
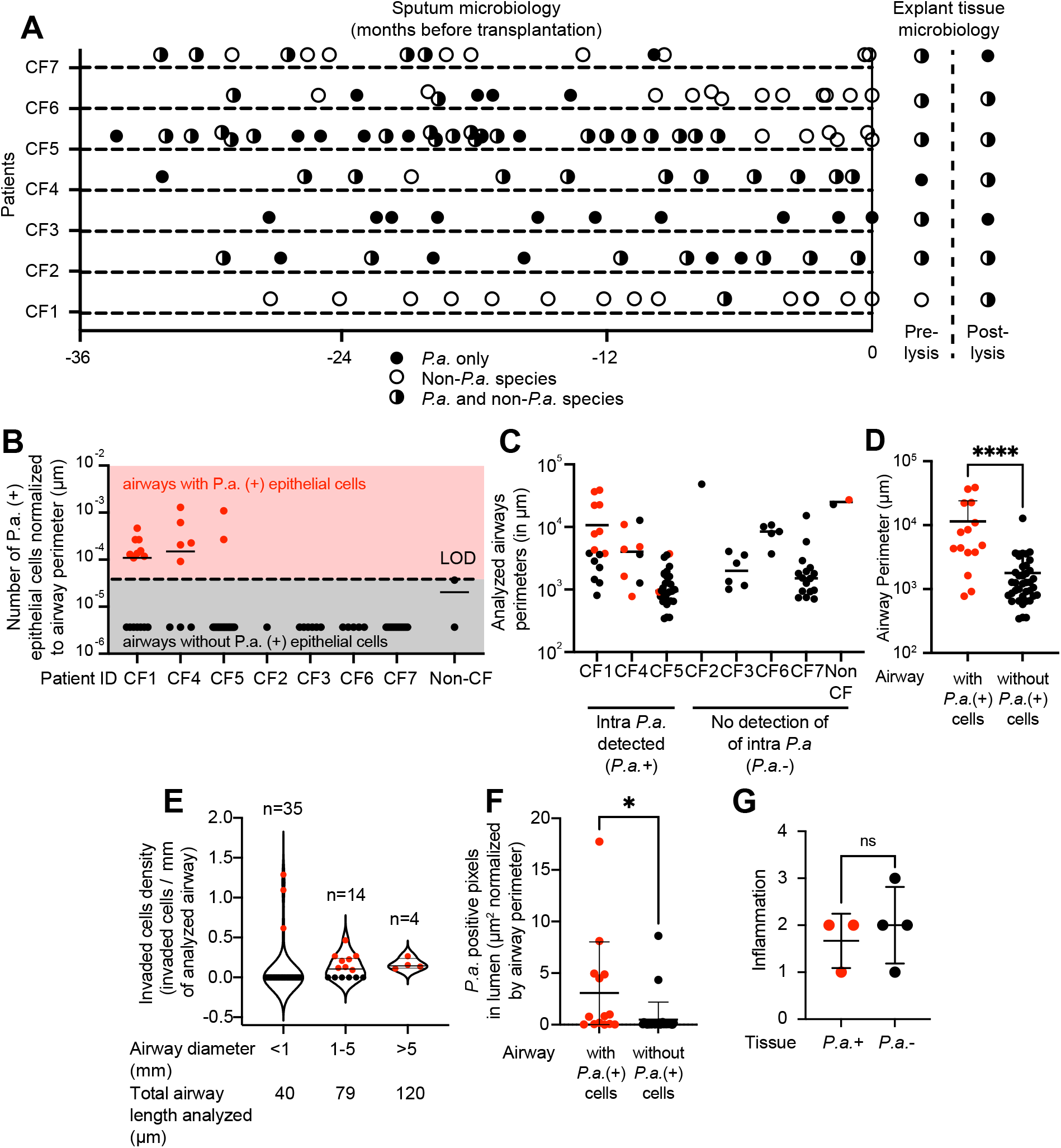
Microbiological, immunohistochemical and histological analysis of a cohort of CF lung explants. (A) Culture-based microbiology of sputum and lung explant tissues from 7 CF patients. For each patient, cultures results are shown for all sputum collected up to 3 years before lung transplantation, as well as pre-lysis and post-lysis lung explant samples. (B) Quantification of airway epithelial cells harboring intracellular *P*.*a*. (referred to as P.a.(+) cells) in airway sections, with the number of *P*.*a*.(+) cells normalized by the perimeter of that airway. LOD limit of detection. The size of airways analyzed are shown according to (C) each patient or (D) the presence or absence of *P*.*a*.(+) cells in that airway. (E) Density of *P*.*a*.(+) cells grouped by the size of their respective airways. For each airway size group, the number of airways and total airway length analyzed are indicated. (F) Quantification of luminal *P*.*a*. in airways with and without *P*.*a*.(+) cells. For B to H, intracellular *P*.*a*. detection by IHC and airway morphometric analyses were performed on thin lung sections stained with anti-*P*.*a*. antibody and hematoxylin. Each dot represents one airway section analyzed, • red for airways with *P*.*a*.(+) cells, • black for airways without *P*.*a*.(+) cells. (G) Histology analysis of H&E stained lung explant samples for regional inflammation. Each dot represents a tissue bloc which are grouped based on tissues that contain airways with and without *P*.*a*.(+) cells.

Next, we performed IHC on lung tissues from all CF (n=7) and non-CF (n=1), for a total of 81 airway sections (perimeter range 343 to 38,827 μm (representative examples shown in Supplemental Figure E3) from 11 tissue blocks. Intracellular *P*.*a*. was detected within airway epithelial cells in tissue sections from CF1, CF4 and CF5, but not the others (Figure 3B). Intracellular *P*.*a*. signals were rare, as they were detected in 15 airways cross-sections, with 1 epithelial cell every 0.8 to 10.9 mm of analyzed airway distance. To characterize the distribution of intracellular *P*.*a*. within the respiratory tract, we measured the airway perimeter of all airway cross-sections analyzed. Our sampling method captured airways in a wide range of sizes, from a perimeter of 343 μm (small bronchioles) to 48,330 μm (bronchi), and this was independent of whether intracellular *P*.*a*. was detected or not (Figure 3C). In patients CF1, CF4 and CF5 where intracellular *P*.*a*. was detected, we compared the size of airways which contained intracellular *P*.*a*. to those that did not from the same patient. This histomorphometric analysis indicated that airways with intracellular *P*.*a*. were 10-fold larger in perimeter than those without intracellular *P*.*a*. (average 11,439 vs. 1,781 μm, p<0.001, Figure 3D). Similarly, when grouping airways of patients CF1, CF4 and CF5 according to their diameters, 12 of 15 airways harboring intracellular *P*.*a*. were bronchioles and bronchi between 1-5 mm or > 5 mm in diameter respectively (Figure 3E). We also analyzed the presence of *P*.*a*. in the airway lumen (Figure 3F) and observed that airways with intracellular *P*.*a*. also displayed 6-fold greater *P*.*a*. signals in the lumen, compared to airways without intracellular *P*.*a*. (3% vs 0.5% all lumen pixels were *P*.*a*. (+)), suggesting a higher luminal bacterial burden. Finally, we asked whether airways associated with intracellular *P*.*a*. were located in lung regions with greater inflammation. We compared histopathological inflammation score of blocs from CF1, CF4 and CF5 to the other patients where no evidence of intracellular *P*.*a*. was detected and did not observe significant differences between the 2 groups (Figure 3F and Supplemental Figure E4).

To better assess the localization of intracellular *P*.*a*. in three dimensions, we performed confocal microscopy of thick tissue sections (1 mm^3^) in a subset of lung explants, using Microbial identification after Passive CLARITY Technique (MiPACT), a tissue clearing technique coupled with immunofluorescence to detect *P*.*a*. and with DAPI and WGA staining to assess the overall tissue structures. Imaging of bronchial structures (Figure 4A) revealed the presence intracellular *P*.*a*. within bronchial epithelial cells (Figure 4B-C) or multinucleated cells (considered polymorphonuclear cell) within the lumen (Figure 4D). We noted that while staining artefacts could be observed in the respiratory zone or the airway lumen when samples were stained with the secondary antibody alone (Figure E5A-D), no intracellular signals were observed in epithelial cells. We also observed the presence of bacterial aggregates in alveoli (Figure E5E-F), compatible with other reports (9). As controls for the specificity of the *P*.*a*. MiPACT-coupled immunofluorescence, we tested a suspension of planktonic *P*.*a*., fixed, hydrogel embedded and processed similarly to lung tissue thick sections. Detection of bacteria, first by immunofluorescence with the polyclonal anti-*P*.*a*. antibody, then by hybridization chain reaction with a *Pseudomonas*-specific probe, confirmed co-localizing *P*.*a*. signals (Figure E5G).

**Figure 4.**
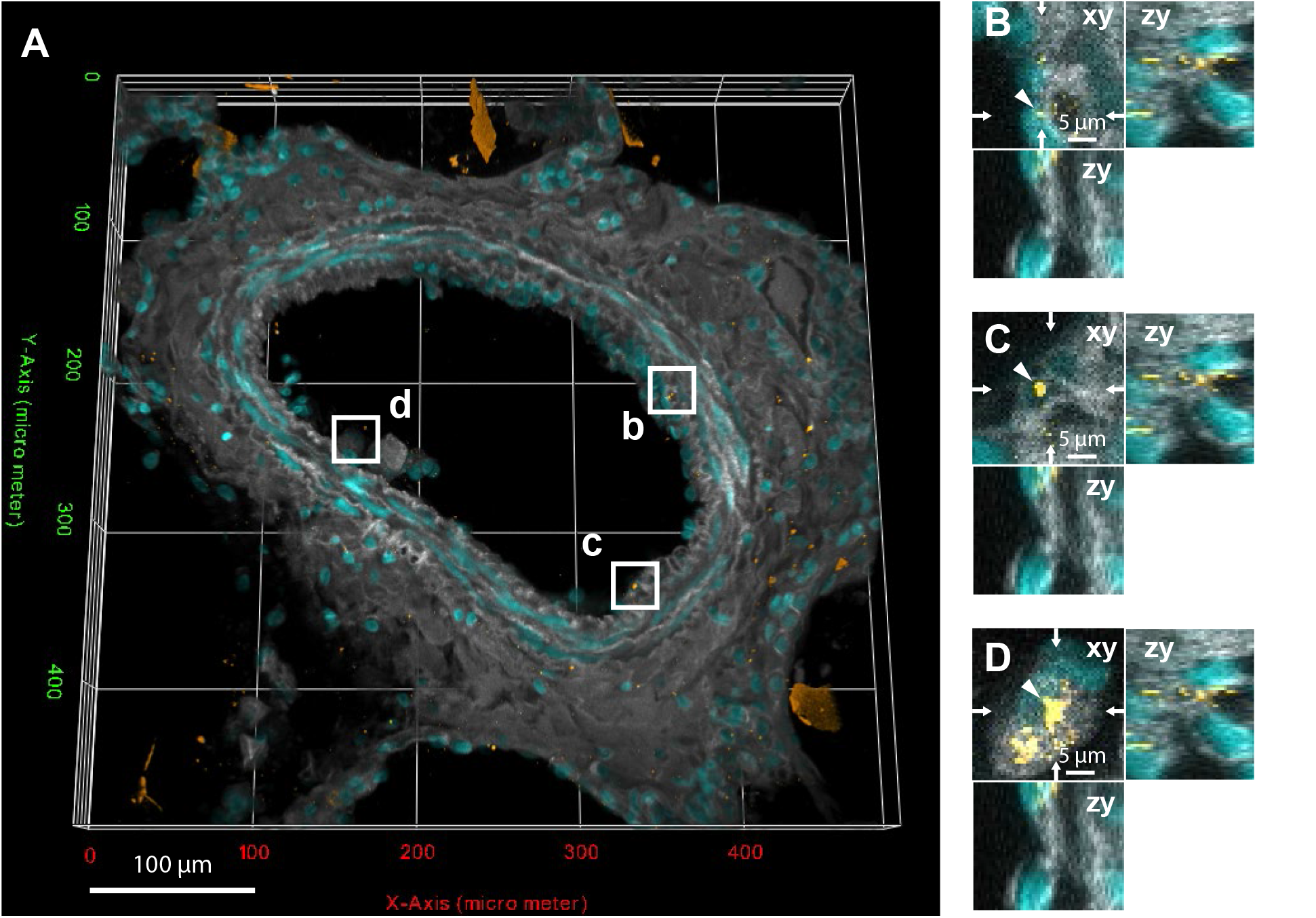
Intracellular *P.a*. detected by 3D MiPACT immunofluorescence analysis of thick lung explant sections. The CF lung explant thick tissue section was cleared and stained with an anti-*P*.*a*. antibody (orange), WGA (grey) and DAPI (cyan) for the lung architecture. (A) Low magnification of a bronchial tissue cross-section. (B-D) High magnification orthogonal projections of areas (white squares) depicting intracellular *P*.*a*.-like signal.

## Discussion

In this manuscript, we first present a case study of a patient with severe CF lung disease undergoing lung transplantation, known for prior *P*.*a*. infection but in whom *P*.*a*. was only recovered once in sputum cultures in the >2 years prior to transplantation. While no extracellular *P*.*a*. was recovered from the lung explant tissues, cultures from lysed tissues indirectly suggested viable intracellular *P*.*a*. Although multiple different cell types, including phagocytes could harbour such intracellular bacterial reservoirs, intracellular *P*.*a*. were detected in airway epithelial cells by immunohistochemistry. We then examined a collection of CF lung explants from patients with a known history of *P*.*a*. infection and found evidence of intracellular *P*.*a*. in the airway epithelium in 3 out of the 7 lungs examined using thin section immunohistochemistry and histology, and thick section immunofluorescence coupled to novel tissue clarification methods. Our findings indicate that intracellular *P*.*a*. are likely relatively rare events most readily detected in lung regions that contain a high bacterial burden and do not correlate with inflammation severity on histopathology. This is the first report of intracellular *P*.*a*. in human CF lung tissues.

Our study has several limitations. First, all of our tissue analyses were performed on lung explant samples. We recognize that end-stage CF lung disease is not representative of most CF individuals with chronic infection, but such samples are the only ones available that provide sufficient tissues for analysis. Furthermore, we sampled the explant lungs in a consistent but random manner and analyzed all airways captured in our samples, but recognize that end-stage CF lung disease displays significant regional heterogeneity. Although we did analyze airways ranging from 0.3 to 48 mm in size, our sampling approach predominantly captured airways in the 10 mm range size. Furthermore, there were significant differences across patients in the total airway perimeters analyzed (from 13 mm to 160 mm), as well as the airway numbers and calibers. For example, large bronchi (diameter >5 mm) were only sampled in 2 patients, and upper lobes were only sampled in 4 patients.

Second, sputum and tissue bacterial cultures may be the gold standard to demonstrate the presence of viable *P*.*a*. but only provide indirect evidence for intracellular *P*.*a*. infection and present several limitations. Sputum cultures may be negative for *P*.*a*. due to incomplete sampling in the setting of low bacterial burden, or due to incomplete bacterial recovery if patients were treated with antibiotics at the time of sputum collection. Sputum may also lack sensitivity for lower respiratory tract infections (33). Finally, our sampling approach and microscopy imaging likely underestimates the prevalence of intracellular *P*.*a*. Conversely, *P*.*a*. recovered from cultures of lysed lung tissues do not necessarily reflect intracellular bacteria. Although we tailored the antibiotic regimens used to kill extracellular bacteria and confirmed that tissue washes post-antibiotics were sterile, it is possible that the tissue lysis procedures also disrupted antibiotic tolerant biofilm aggregates (34), and that bacteria recovered post-lysis were in fact extracellular bacteria that survived antibiotic treatment. Finally, we used non-CF tissue section as negative controls, as there were no CF patients for which sputum and lung explant tissue cultures were both negative for *P*.*a*.

Several previous microscopy studies have detected *P*.*a*. in the airways of CF lung tissues, either by analyzing lung explants as we have, or post-mortem autopsy tissues (9–12, 30, 32). Consistent with other studies, we did note the presence of *P*.*a*. microcolonies or leukocyte-trapped bacteria in the mucopurulent material filling the airway lumen and did not observe significant bacteria adherent to the epithelial surfaces. While most studies never reported evidence of bacterial invasion within CF lung tissues (9, 10, 32), only one study performed a thorough analysis of airway lumen and epithelia (12). Most importantly, no study specifically sought for evidence of intracellular *P*.*a*. Our microscopy analyses used polyclonal *P*.*a*.-specific antibodies that were able to detect clinical isolates and were optimized to detect intracellular *P*.*a*. signals, typically micrometer signals rather than microcolonies (> 100 μm diameter) found within the airway lumen. While we did find evidence of luminal and extracellular bacteria, this bacterial burden was relatively low, likely as a result of our tissue sampling and processing procedures that wash out luminal secretions.

The ability of bacteria classically known as extracellular pathogens to reside, and in some cases to replicate intracellularly, is likely an underappreciated mechanism underlying recurrent and chronic bacterial infections, such as chronic urinary tract infections, endocarditis or osteomyelitis (35–37). Among organisms causing chronic CF infection or colonization, *Burkholderia cepacia* complex (Bcc), *Haemophilus influenzae* or *Staphylococcus aureus* have also been reported to invade respiratory epithelial cells *in vitro*. These internalization mechanisms have been reviewed extensively in (38–40). Histology-based studies of CF patient lung tissues or *ex vivo* infection experiments remain scarce but show diverse patterns of intracellular localization in the CF airway that are dependent on bacterial genomic or microbe-microbe interactions. For example, the invasive *Burkholderia cenocepacia* ET12 strain has been observed intracellularly and localized diffusively along airways and parenchymal tissues of CF lungs (41), while other *B. cenocepacia* complex strains have been primarily found within phagocytic cells in the mucus of large airways (11).

Several studies have examined intracellular bacteria in tissues of the respiratory tract, although not from individuals with CF. *S. aureus* is the most prevalent pathogen colonizing the respiratory tract of young individuals with CF (42). While *S. aureus* clearly resides in the mucus of obstructed airways (43, 44), and are found as bacterial aggregates in CF sputum (31), it also colonizes the CF sinonasal passages (45). Interestingly, *S. aureus* can invade different cell types, including epithelial and endothelial cells (39, 46) and its intracellular form has been associated with infections often refractory to antibiotic treatment. Notably, intracellular *S. aureus* has been associated with recurrent rhinosinusitis and has been detected *in situ* in endonasal tissues including epithelial cells (36, 47, 48).

In our study, we performed IHC, a well-established microscopy method using a well validated polyclonal antibody. While such analyses do not reveal whether the detected bacteria are viable or not, they are highly sensitive and allowed us to focus on detection of intracellular bacteria within the epithelium. The use of probes that could confirm metabolic activity in intracellular bacteria would be of value. We also employed MiPACT (passive CLARITY technique), a recently developed approach which can reveal the 3D spatial distribution of CF microbes *in situ* and their relationship with their host cells and tissue structures by confocal microscopy (49, 50). Recent advances of this technology allowed the study of the biogeography of CF pathogen in sputum or the upper airways with resolution at the species level using multiplexed molecular probes (31, 51). Here, MiPACT was applied for the first time to human lung tissues to visualize CF airways, and combined with immunofluorescence, revealed rare events of intracellular bacteria within the airway epithelium. Future studies of MiPACT in combination with metabolic reporter (31, 52) may provide further evidence of metabolically active and thus viable intracellular bacterial.

In conclusion, we reveal for the first time evidence for intracellular *P*.*a*. in the airway epithelium of CF lung explant tissues. These findings highlight the plausible occurrence of *P*.*a*. intracellular lifestyle in CF lung infection. Further studies are warranted to understand the contribution of *P*.*a*. intracellular lifecycle to the pathogenesis and persistence of CF lung infections.

## Supporting information

Supplementary information Malet et al

## Acknowledgements

We would like to acknowledge the CR CHUM Biobank for access to tissues and clinical data, the Cystic Fibrosis Canada registry for clinical data. We would like to thank the patients for participating to this study. We thank the RI MUHC Histopathology platform for their technical assistance with the IHC.

