## Supplementary information Malet et al for "Intracellular *Pseudomonas aeruginosa* within the airway epithelium of Cystic Fibrosis lung tissues"

Short running title: Intracellular *Pseudomonas aeruginosa* in CF lung tissues

Karim Malet (1), Emmanuel Faure (2,3), Damien Adam (4,5), Jannik Donner (1), Lin Liu (6, 7), Sarah-Jeanne Pilon (8), Richard Fraser (8), Peter Jorth (9-12), Dianne K. Newman (12,13), Emmanuelle Brochiero (4,5), Simon Rousseau (1), Dao Nguyen (1,14)

**Affiliations**
[1]Meakins-Christie Laboratories, Research Institute of the McGill University Health Centre (RI-MUHC), Montreal, Canada
[2]Univ. Lille, CNRS, Inserm, CHU Lille, Institut Pasteur Lille, U1019 -UMR 9017 -CIIL -Center for Infection and Immunity of Lille, F-59000 Lille, France
[3]CHU Lille, Service Universitaire de Maladies Infectieuses, F-59000 Lille, France
[4]Centre de recherche du Centre Hospitalier de l'Université de Montréal (CR-CHUM), Montreal, Canada
[5]Département de Médecine, Université de Montréal, Montréal, QC, Canada.
[6]Department of Respiratory and Critical Care Medicine, Guizhou Provincial People's Hospital, Guiyang, Guizhou 550002, China.
[7]NHC Key Laboratory of Pulmonary Immunological Diseases, Guizhou Provincial People's Hospital, Guiyang, Guizhou 550002, China.
[8]Department of Pathology, McGill University Health Centre (MUHC), Montreal, Canada
[9]Cedars-Sinai Medical Center, Department of Pathology and Laboratory Medicine, Los Angeles, California, USA.
[10]Cedars-Sinai Medical Center, Department of Medicine, Los Angeles, California, USA.
[11]Cedars-Sinai Medical Center, Department of Biomedical Sciences, Los Angeles, California, USA.
[12]Division of Biology and Biological Engineering, California Institute of Technology, Pasadena, California, USA.
[13]Division of Geological and Planetary Sciences, California Institute of Technology, Pasadena, California, USA.
[14]Department of Medicine, McGill University, Montreal, Canada

Corresponding author: Dao Nguyen
Research Institute of the McGill University Health Centre
1001 Decarie Blvd, Mailstop EM3.2219
Montreal, QC, H4A 3J1, Canada


### Supplemental Material and Methods

*Culture-based microbiological analysis of lung tissues*

Fresh lung tissue samples were weighed, placed in 1 mL of sterile PBS, vortexed and incubated for 1 h at 4°C to recover extracellular bacteria. Viable extracellular bacteria (pre-lysis culture) were counted by plating serial dilutions of the PBS supernatant on LB agar (Difco 244620), Pseudomonas isolation agar (PIA, BD #292710), Blood agar (BA, BD #254071), Mannitol salt agar (MSA, BD 211407) and McConkey agar (MCA, BD 299769) plates. Tissues were subsequently washed three times in sterile PBS and mixed with an antibiotic cocktail by vortexing to kill extracellular bacteria, particularly *P.a.* The antibiotic cocktail consisted of 100 µg/mL tobramycin (Sigma-Aldrich) and/or 20 µg/mL colistin (MP Biomedicals) and was selected based on the antibiotic susceptibility of *P.a.* from each patient's most recent sputum culture. After 30 min incubation at room temperature, the antibiotic-containing buffer was discarded, and tissues were washed twice with sterile PBS to remove any residual antibiotic. The last wash solution was plated on LB agar as control to ensure sterilization. Tissues were then lysed in 1 mL sterile PBS containing 0.5% Triton (Sigma-Aldrich) using Teflon beads (0.9 – 2.0 mm, Next Advance #SSB14B) and tissue lyser (Bullet Blender, Next Advance) (5 min x 2 cycles). The viable bacteria recovered from the lysed tissues (referred to as post-lysis cultures) were counted as colony forming units (CFU) with serial dilution of the tissue lysates and plating on LB, PIA, MSA, MCA and BA plates. All morphologically distinct bacterial clones isolated from those plates were analyzed by Matrix Assisted Laser Desorption/Ionization - Time of Flight (MALDI-TOF) (Bruker UltrafleXtreme MALDI Biotyper RTC, Bruker Daltonics) for specie identification using standard protocols.

*Histology and immunohistochemistry (IHC) detection of P. aeruginosa*

Fresh lung tissues were fixed in 10% formalin and embedded in paraffin embedding for thin sectioning (0.4 mm). For IHC, the thin sections were stained with hematoxylin, 1:3000 rabbit anti-Pseudomonas polyclonal antibody (Abcam, ab68538) at 37°C during 2 h, followed by anti-rabbit-HRP conjugated secondary antibody (DISCOVERY OmniMap anti-Rb, #760-4311, Roche Diagnostics) and the chromogenic reagent (Discovery Purple kit #760-229, Roche Diagnostics) using an automated slide preparation system (Ventana Discovery Ultra, Roche Diagnostics). For histopathology scoring, Hematoxylin and Eosin stained thin sections were analyzed in a blinded manner by two clinical pathologists using a semi-quantitative score to assess the severity of inflammation and pathology in the airway and parenchymal tissues sections.

*Histology image acquisition and analyses*

Stained lung thin sections were scanned at 20x and 40x using an Aperio AT Turbo scanning system (Leica), creating pyramid tiled svs files analyzed using the Qpath software version 0.3.2. First, airway structures dominated by ciliated epithelial cells were identified by the hematoxylin staining. The brush tool was used to manually draw two Regions Of Interest (ROI), one that encompasses the entire airway (airway ROI) up to the smooth muscle layer if present, and one that defines the lumen of the airway (lumen ROI). The airway ROI was used for assessment of the airway perimeter (in µm) and was further divided into smaller areas for manual assessment of *P.a.* by immunohistochemistry. Positive signals for intracellular *P.a.* were defined as 0.5-2 µm-sized purple rod-to-sphere-like signals within the boundaries of airway epithelial cells. The number of infected epithelial cells was reported for each airway and normalized by the airway perimeter. For each airway, quantification of *P.a.* within the airway lumen was performed by measuring the total

*P.a*-positive signal in the lumen ROI and normalizing it by the airway perimeter. All images were randomized and analyzed in a blinded manner.

*Lung tissue clearing by passive CLARITY for MiPACT-IF analysis*
Tissues were cleared as previously done in (DePas mBio 2016). Briefly, fresh lung tissues were fixed in 4% PFA for 2 h at RT with shaking. Tissues were then kept at 4°C for 24 h, washed twice with PBS and (pH=7.4), immersed in cold B4P1 hydrogel solution (4% of acrylamide/bis solution 29:1 (Biorad, 1610156), 1% PFA, 0.25% of VA044 thermoinitiator (Sigma-Aldrich, 2997-92-4)) and incubated overnight at 4°C with shaking. The B4P1 solution was then discarded, fresh ice-cold B4P1 was added and allowed to polymerize during 3 to 4 h at 37°C under oxygen-free conditions. Hydrogel-embedded tissues were sectioned into thick sections (1 mm), washed twice in PBS and incubated in 8% Sodium dodecyl sulfate (SDS) PBS solution (pH=7.4) at 37°C with shaking at 100 r.p.m during 5 to 6 weeks for clearing of lung tissues. Following clearing, lung tissues were washed three times in sterile PBS (pH=7.4) prior to staining.

*MiPACT-Immunofluorescence (IF) for P. aeruginosa detection*
Cleared lung tissue thick sections were first stained for *P.a.* using a 1:400 mouse anti-*P.a.* monoclonal (Thermo Fisher, # MA183430), diluted in PBS supplemented with 2.5% goat serum, by incubating in staining solution for 1 day at RT with gentle shaking, followed by sterile PBS washes x3 for 2h each and a final wash of 16h. Secondary antibody staining was performed using 1:400 goat anti-mouse Alexa Fluor 555 (Thermo Fisher, # A32727) antibody diluted in PBS supplemented with 2.5% goat serum during 3 days at RT, followed by sterile PBS washes x3 for 2 h each and a final wash of 16 h. Finally, tissues were stained with Alexa-488 conjugated wheat-germ agglutinin (WGA, Thermo Fisher, #W11261) (20µg/mL) in 1 mL PBS (Wisent, 311-513-CL) for 16 h, washed in OBS for 16 h at RT in the dark. Samples were finally cleared for 24 h in Refracting Index Matching Solution (RIMS) : 80% HistoDenz (w/v) (Sigma, #D2158) in 20 mM phosphate buffer with 0.1% Tween 20, 0.01% sodium azide pH 7.5, 10 µg/ml DAPI.

*Bacterial strains, culture conditions and processing of planktonic bacteria for immunofluorescence (IF) and immunohistochemistry (IHC) analysis*
All bacterial strains used in this study are summarized in Supplemental Table E2. Strains were streaked from frozen stocks onto LB agar (Wisent), incubated overnight at 37˚C, and isolated colonies were picked to inoculate 5 mL of liquid LB medium. After overnight incubation at 37°C with shaking, bacterial cells were pelleted, washed twice with sterile phosphate buffered saline (PBS), and resuspended in LB medium (~10^8 CFU/mL). For IF, 500 µL of the bacterial suspension was seeded in 24-well plate containing uncoated sterile 15mm round borosilicate glass coverslips (0.13 to 0.17 mm thick, Fisherbrand), centrifuged at 1500 r.p.m for 5 min and incubated for 2 h at 37˚C. Coverslips were then washed in PBS, fixed in 4% PFA for 15 min, washed again in PBS and bacteria were stained for 16 h at 4 ˚C using a rabbit anti-Pseudomonas polyclonal antibody (Abcam, #ab68358) diluted in PBS containing 1% BSA (Sigma-Aldrich) and 2% goat serum (Sigma-Aldrich). Bacteria were then washed and stained for 1 h at RT with a Rhodamine Red-X donkey anti-rabbit antibody (Jackson ImmunoResearch, #711-295-152) in staining buffer containing DAPI (10 µg/mL). Coverslips were finally washed and mounted in Fluoromount-G (Thermo Fisher Scientific). For IHC, ~5x10^8 cells were pelleted and fixed in 4% PFA for 15 min before dehydration and paraffin embedding. Thin sections were processed as done with lung tissue samples.

*Bacterial detection by MiPACT-hybridization chain reaction (HCR)*

Bacteria were grown overnight in LB medium at 37°C with shaking, then fixed in 4% PFA for 16 h at 4°C. Fixed bacterial cells were concentrated to $10^{11}$ cells/mL and incubated for 16 h at 4°C in fresh filter-sterilized 4% (vol/vol) 29:1 acrylamide:bis-acrylamide and 0.25% (wt/vol) VA-044 hardener in PBS. Samples were transferred to an anaerobic chamber for 15 min to remove oxygen from the tube headspace, then incubated in a 37°C water bath for 3 h to harden samples. Samples were then cleared in SDS (see below passive CLARITY technique) and processed for HCR. 1 $mm^3$ cleared blocks of planktonically grown *P.a.* PA14 were placed in 500 ml 1 mg/ml lysozyme + 0.05 mg/ml lysostaphin buffer and incubated for 1 h at 37°C to break down peptidoglycan cell walls. Blocks were transferred to 15 ml PBS and incubated rocking at 25°C for 60m. Samples were transferred to fresh PBS for an additional 60 min wash. 5 ml of each HCR probe, EUB338 or NON338 (negative control), was diluted into filter sterilized 500 ml HCR hybridization solution (300 ml 20X SSC, 600 ml Formamide, 300 mg dextran sulfate, with water up to 3 ml total volume). Washed blocks were transferred to appropriate HCR solutions and incubated shaking at 300 rpm at 46°C in shaking heat block overnight. Blocks were transferred to 50 ml of 84 mM NaCl FISH wash buffer and incubated for 6 h at 52°C in a water bath. 22 ml each of 3 mM B1H1 and B1H2 (Alexa Fluor 647, Molecular Technologies) HCR hairpin probes were heat shocked for 90s at 95°C in a thermal cycler and incubated in the dark at 25°C for 30 m. Washed blocks and 5 ml of each denatured HCR probe were transferred to 110 ml HCR amplification buffer (200 mg dextran sulfate, 200 ml 20X SSC, with water up to 2 ml). Incubate gently shaking overnight at room temperature in the dark by covering with foil. Blocks were transferred to 50 ml 337.5 mM NaCl FISH wash buffer and incubated 3 h at 48°C in a water bath.

*Confocal microscopy*

Stained lung tissue and bacterial sections were mounted on perfusion chambers (Electron Microscopy Services) in RIMS and imaged by confocal microscopy (Zeiss LSM 800) using the 405nm, 488nm, 561nm laser channels. The following objectives were used: EC-Plan Neofluar 10X/0.30-numerical aperture, Plan-Apochromat 10x/0.45-numerical aperture M27 with working distance 2 mm, or LD LCI Plan-Apochromat 25x/0.8-numerical aperture Imm Corr DIC M27 multi-immersion objective with a 0.57 mm working distance, using glycerol for immersion. Z-stacks were collected in 12-bit mode with 2-line averaging at 1024x1024 resolution. Bacterial suspensions on coverslip were imaged with a Zeiss LSM700 confocal microscope equipped with 405 nm and 543 nm lasers and a plan-apochromat 63x/1.40 oil DIC M27 immersion objective. Images were exported as czi files and analyzed and processed using ImageJ (Version 2.0) for single plane, maximum intensity projection, orthoview image processing or 3D rendering video, and using ICY (version 2.4.2.0) for 3D reconstructions.

**Table E1. Patient characteristics and sputum analysis**

| CF ID | CF1 | CF2 | CF3 | CF4 |
|---|---|---|---|---|
| Age | 48 | 22 | 48 | 30 |
| Sex | F | M | F | F |
| CFTR variant | F508del/711+1GT | 1525-1G>A/ 1525-1G>A | F508del/F508del | F508del/F508del |

**CF1 – Microbiology**

| DoA | DBT | *S.m.* | *A.f.* | Other |
|---|---|---|---|---|
| 17-05-23 | 0,00 | X | | *B.v.* |
| 17-04-20 | -33,00 | X | X | |
| 17-02-28 | -84,00 | X | X | |
| 17-03-01 | -83,00 | X | X | |
| 17-01-31 | -112,00 | X | X | |
| 16-11-01 | -203 | X | X | *P.a.* |
| 16-08-02 | -294 | X | X | |
| 16-06-30 | -327 | X | X | |
| 16-05-19 | -369 | X | X | F/Y |
| 16-03-03 | -446,00 | X | X | |
| 15-12-17 | -523 | X | X | |
| 15-10-22 | -579 | X | X | |
| 15-08-27 | -635 | X | X | |
| 15-05-21 | -733 | X | X | |
| 15-02-15 | -828 | X | X | |
| 13-01-01 | -1603 | | | S.a. |

**CF2 – Microbiology**

| DoA | DBT | *P.a.* | *S.m.* | Other |
|---|---|---|---|---|
| 17-07-29 | 0 | | | |
| 17-07-30 | 1,00 | X | X | |
| 17-07-10 | -19,00 | X | | NOF |
| 17-05-05 | -85,00 | X | | NOF |
| 17-03-02 | -149,00 | X | X | NOF |
| 17-01-30 | -180,00 | X | | NOF |
| 16-12-21 | -220,00 | X | | NOF |
| 16-11-16 | -255,00 | X | X | NOF |
| 16-08-17 | -346,00 | X | | H.i., NOF |
| 16-04-06 | -479,00 | X | | NOF |
| 15-12-03 | -604,00 | X | | |
| 15-09-09 | -689 | X | | F/Y, NOF, *C.a.* |
| 15-05-07 | -814 | X | | NOF |
| 15-02-17 | -893 | X | | NOF, *S.a.* |

**CF3 – Microbiology**

| DoA | DBT | *P.a.* | Other |
|---|---|---|---|
| 17-10-29 | 0,00 | X | |
| 17-09-13 | -46,00 | X | |
| 17-06-29 | -122,00 | X | |
| 17-01-12 | -290,00 | X | NOF |
| 16-10-13 | -381,00 | X | NOF |
| 16-07-26 | -460,00 | X | |
| 16-03-10 | -598,00 | X | NOF |
| 16-01-07 | -661 | X | NOF |
| 15-12-17 | -682 | X | NOF |
| 15-07-22 | -830 | X | NOF |
| 15-01-22 | -1011 | X | NOF |

**CF4 – Microbiology**

| DoA | DBT | *P.a.* | *A.x.* | *S.m.* | *A.f.* | Other |
|---|---|---|---|---|---|---|
| 18-01-10 | 0,00 | X | | X | X | NOF, *C.i., S.mu.* |
| 17-12-19 | -22,00 | X | X | | | |
| 17-10-27 | -75,00 | X | | X | | NOF, *C.i.* |
| 17-08-28 | -135,00 | X | | X | | NOF, C.i., S.mu. |
| 17-06-16 | -208,00 | X | X | X | | NOF, *F.m.* |
| 17-04-28 | -257,00 | X | | X | | NOF, *F.mu.* |
| 16-12-14 | -392,00 | X | | | | *H.i.* |
| 16-09-16 | -481,00 | X | | X | X | NOF |
| 16-05-13 | -607,00 | | | X | X | NOF |
| 16-02-26 | -684,00 | X | | | X | NOF, *C.i.* |
| 15-12-18 | -754,00 | X | | | | NOF |
| 15-06-05 | -950,00 | X | | | | NOF |
| 15-02-09 | -1066,00 | X | | X | | F/Y, NOF |

Definition of abbreviations: DoA=Date of Analysis, DBT=Days before transplantation, *P.a.=Pseudomonas aeruginosa, B.c.=Burkholderia cepacia, S.a.=Staphylococcus aureus, Achromobacter, A.b.=Acetinobacter baumanii, S.m.=Stenotrophomonas maltophilia, B.v.=Brevundimonas vesicularis, E.a.=Enterobacter aerogenes, C.a.=Candida albicans, A.f.=Aspergillus fumigatus,* NOF= normal oropharyngeal flora, F/Y=Fungal/Yeast infection, *C.i.=Chryseobacterium Indologenes, F.m.=Flavobacterium meningosepticum, S.mu=Sphingobacterium multivorum, H.i.= Hemophilus influenzae, B.a.=Burkholderia anthina, P.ag.=Pantoea agglomerans, P.d.=Pseudomonas denitrificans*

**Table E1. Patient characteristics and sputum microbiology (continued)**

| CF ID | CF5 | | | | | |
|---|---|---|---|---|---|---|
| Age | 22 | | | | | |
| Sex | M | | | | | |
| CFTR variant | F508del/F508del | | | | | |
| **Microbiology** | | | | | | |
| DoA | DBT | *P.a.* | *B.c.* | *S.a.* | *A.x.* | *Other* |
| 18-03-01 | 0 | | X | | | |
| 18-02-23 | -6 | | X | X | | B.a. |
| 18-01-12 | -48 | | X | X | | B.a. |
| 18-01-01 | -59 | | X | X | | |
| 17-12-01 | -90 | | X | X | | B.a. |
| 17-10-01 | -151 | | X | X | | B.a. |
| 17-08-01 | -212 | X | | X | | |
| 17-07-01 | -243 | X | X | X | | |
| 17-06-09 | -265 | X | X | X | | |
| 17-05-01 | -304 | X | X | X | | |
| 17-03-31 | -335 | X | X | X | | |
| 17-03-01 | -365 | X | X | | | |
| 17-02-03 | -391 | X | X | X | | |
| 16-11-01 | -485 | X | | | | |
| 16-10-01 | -516 | X | X | | | |
| 16-09-09 | -538 | X | | X | | NOF |
| 16-09-01 | -546 | X | | X | | |
| 16-08-26 | -552 | X | | X | | |
| 16-08-01 | -577 | X | | X | | |
| 16-07-08 | -601 | X | X | X | | |
| 16-07-01 | -608 | X | X | X | | |
| 16-06-01 | -638 | X | | | | |
| 16-05-01 | -669 | X | | X | | |
| 16-04-01 | -699 | X | | | | |
| 16-02-01 | -759 | X | | | | |
| 16-01-01 | -790 | X | | | | |
| 15-11-01 | -851 | X | X | X | | |
| 15-10-01 | -882 | X | X | X | | |
| 15-09-18 | -895 | X | | X | | |
| 15-08-07 | -937 | X | X | X | | NOF |
| 15-07-03 | -972 | X | | X | | |
| 15-04-26 | -1040 | X | | | | |
| 15-03-22 | -1075 | X | X | | | F/Y |
| 15-03-01 | -1096 | X | X | | | |

| CF ID | CF6 | | | |
|---|---|---|---|---|
| Age | 26 | | | |
| Sex | M | | | |
| CFTR variant | F508del/F508del | | | |
| **Microbiology** | | | | |
| DoA | DBT | *P.a.* | *A.x.* | Other |
| 18-04-08 | 0 | | X | |
| 18-03-09 | -30 | | X | NOF |
| 18-02-04 | -63 | | X | NOF |
| 18-01-31 | -67 | | X | S.a., NOF |
| 17-12-06 | -123 | | X | |
| 17-11-08 | -151 | | X | NOF |
| 17-09-13 | -207 | | X | *P.d.*, NOF |
| 17-08-30 | -221 | | X | NOF |
| 17-08-04 | -247 | | X | NOF, A.b., H.i. |
| 17-06-14 | -298 | | X | *F.m., P.ag.,S.m.,* NOF |
| 17-02-17 | -415 | X | | NOF |
| 16-11-02 | -522 | X | | NOF |
| 16-10-12 | -543 | X | | NOF |
| 16-08-19 | -597 | X | X | NOF |
| 16-08-05 | -611 | | X | NOF |
| 16-04-29 | -709 | X | | NOF |
| 16-03-07 | -762 | | X | *P.d.*, NOF |
| 15-11-11 | -879 | X | X | NOF |
| 15-03-18 | -1117 | X | | NOF |
| 15-01-28 | -1166 | X | | NOF |

| CF ID | CF7 | | | | | | |
|---|---|---|---|---|---|---|---|
| Age | 15 | | | | | | |
| Sex | F | | | | | | |
| CFTR variant | DF508/G542X | | | | | | |
| **Microbiology** | | | | | | | |
| DoA | DBT | *P.a.* | *S.a.* | *A.x.* | *C.a.* | *A.f.* | Other |
| 18-05-07 | 0 | | | | | | |
| 18-05-03 | -4 | | | | | | *E.a.* |
| 18-04-27 | -10 | | | | | | F/Y |
| 17-07-25 | -286 | | | | X | X | F/Y |
| 17-07-11 | -300 | X | | | | | NOF |
| 17-04-04 | -398 | | | X | X | X | F/Y |
| 16-11-01 | -552 | | X | X | | X | |
| 16-09-28 | -586 | | | X | | | F/Y |
| 16-08-30 | -615 | X | | X | | | F/Y |
| 16-08-05 | -640 | X | | X | | | F/Y |
| 16-04-20 | -747 | | | X | | X | F/Y |
| 16-03-21 | -777 | | | | | X | F/Y |
| 16-02-23 | -804 | X | | X | | X | |
| 15-12-08 | -881 | | | X | | X | |
| 15-10-20 | -930 | X | | X | | X | |
| 15-09-01 | -979 | X | | X | | X | |
| 15-08-04 | -1007 | | | X | | X | |
| 15-04-14 | -1119 | X | X | X | | X | |
| 15-03-17 | -1147 | | | X | | X | |
| 15-02-10 | -1182 | | | | | | NOF |
| 15-01-13 | -1210 | X | | X | | X | F/Y |

| | | | | | |
|---|---|---|---|---|---|
| 14-11-01 | -1216 | X | X | X | X |

**Table E2. List of strains and clinical isolates used in this study**

| Strain / Isolate | Reference | DN# | Patient ID | Pre-/Post-lysis isolate* | Region** | *MALDI analysis (Biotyping)* |
|---|---|---|---|---|---|---|
| PAO1 | Jacobs et al. 2003 | DN145 | N.A. | N.A. | N.A. | N.A. |
| *Burkholderia cenopacia* | Jane Burns | CS270 | N.A. | N.A. | N.A. | N.A. |
| *Proteus mirabilis* | ESUT #698B Enugu State University nigeria | DN836 | N.A. | N.A. | N.A. | N.A. |
| *Stenotrophomonas maltophilia* | Jane Burns | CS292 | N.A. | N.A. | N.A. | N.A. |
| 8 | This study | N.A. | CF1 | Post (Intracellular) | RM2 | *P. aeruginosa* |
| 10 | This study | N.A. | CF1 | Post (Intracellular) | RL2 | *P. aeruginosa* |
| 104 | This study | N.A. | CF1 | Post (Intracellular) | RM1 | *P. aeruginosa* |
| 105 | This study | N.A. | CF1 | Post (Intracellular) | RM2 | *P. aeruginosa* |
| 106 | This study | N.A. | CF1 | Post (Intracellular) | RL1 | *P. aeruginosa* |
| 39 | This study | N.A. | CF2 | Pre (Extracellular) | RU1 | *S. aureus* |
| 42 | This study | N.A. | CF2 | Post (Intracellular) | RM1 | *P. aeruginosa* |
| 43 | This study | N.A. | CF2 | Post (Intracellular) | RM2 | *P. aeruginosa* |
| 44 | This study | N.A. | CF2 | Post (Intracellular) | RM1 | *P. aeruginosa* |
| 45 | This study | N.A. | CF2 | Post (Intracellular) | RL1 | *P. aeruginosa* |
| 47 | This study | N.A. | CF2 | Pre (Extracellular) | RU1 | *P. aeruginosa* |
| 48 | This study | N.A. | CF2 | Post (Intracellular) | RU1 | *P. aeruginosa* |
| 65 | This study | N.A. | CF3 | Post (Intracellular) | RU1 | *P. aeruginosa* |
| 66 | This study | N.A. | CF3 | Post (Intracellular) | RU2 | *P. aeruginosa* |
| 67 | This study | N.A. | CF3 | Post (Intracellular) | RL1 | *P. aeruginosa* |
| 68 | This study | N.A. | CF3 | Pre (Extracellular) | RU1 | *P. aeruginosa* |
| 69 | This study | N.A. | CF3 | Pre (Extracellular) | RM1 | *P. aeruginosa* |
| 70 | This study | N.A. | CF3 | Pre (Extracellular) | RL1 | *P. aeruginosa* |
| 71a | This study | N.A. | CF4 | Pre (Extracellular) | RU2 | *P. aeruginosa* |
| 71b | This study | N.A. | CF4 | Pre (Extracellular) | RU2 | *Achromobacter ruhlandii* |
| 72 | This study | N.A. | CF4 | Pre (Extracellular) | LU3 | *P. aeruginosa* |

| | | | | | | |
|---|---|---|---|---|---|---|
| 73 | This study | N.A. | CF4 | Pre (Extracellular) | LU2 | *P. aeruginosa* |
| 74 | This study | N.A. | CF4 | Post (Intracellular) | RU2 | *P. aeruginosa* |
| 75 | This study | N.A. | CF4 | Post (Intracellular) | RL2 | *P. aeruginosa* |
| 76 | This study | N.A. | CF4 | Post (Intracellular) | RL3 | *P. aeruginosa* |
| 77a | This study | N.A. | CF5 | Pre (Extracellular) | RU1 | *P. aeruginosa* |
| 77b | This study | N.A. | CF5 | Pre (Extracellular) | RU1 | *Burkholderia anthina* |
| 78 | This study | N.A. | CF5 | Pre (Extracellular) | RM1 | *P. aeruginosa* |
| 79 | This study | N.A. | CF5 | Pre (Extracellular) | RL1 | *P. aeruginosa* |
| 80 | This study | N.A. | CF5 | Post (Intracellular) | RU1 | *P. aeruginosa* |
| 82 | This study | N.A. | CF5 | Post (Intracellular) | RL1 | *P. aeruginosa* |
| 92 | This study | N.A. | CF6 | Pre (Extracellular) | RU1 | *P. aeruginosa* |
| 93 | This study | N.A. | CF6 | Pre (Extracellular) | RL1 | *P. aeruginosa* |
| 94a | This study | N.A. | CF6 | Post (Intracellular) | RU1 | *P. aeruginosa* |
| 94b | This study | N.A. | CF6 | Post (Intracellular) | RU1 | *Achromobacter ruhlandii* |
| 95 | This study | N.A. | CF6 | Post (Intracellular) | RL1 | *P. aeruginosa* |
| 96 | This study | N.A. | CF7 | Post (Intracellular) | RL1 | *P. aeruginosa* |
| 98 | This study | N.A. | CF7 | Post (Intracellular) | RL1 | *P. aeruginosa* |
| 99 | This study | N.A. | CF7 | Pre (Extracellular) | RL1 | *Staphylococcus warneri* |
| 100 | This study | N.A. | CF7 | Post (Intracellular) | RM2 | *P. aeruginosa* |

*Pre-lysis (or extracellular) isolates were retrieved from tissue after resuspension in PBS and before antibiotic treatment and lysis. Post-lysis (or intracellular) isolates were retrieved from tissue after antibiotic treatment and lysis.

**Lung region from which isolate was retrieved (*e.g.* LU1 = Left Upper lobe sample 1 ; RM2 = Right Middle lobe sample 2 ; RL1 = Right Lower lobe sample 1 ; *etc.*)

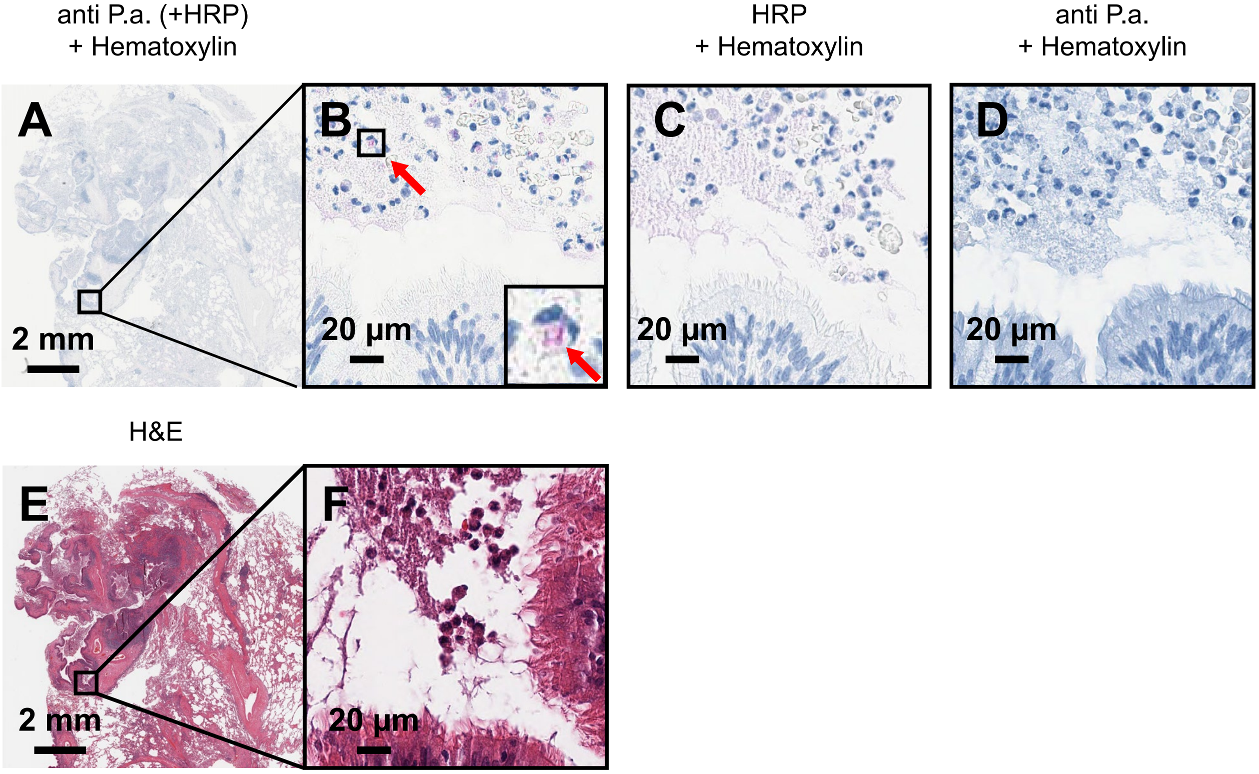

**Supplementary figure E1. Staining controls for IHC analysis.** CF lung explant thin section was analyzed for the following control staining (A) Hematoxylin, anti-*P.a.* primary antibody and secondary anti-rabbit antibody coupled with Horseradish Peroxidase (HRP). (B) High magnification of (A), with inset showing intracellular *P.a.* signal in luminal polymorphonuclear cell (indicated by black arrow). (C) Hematoxylin, HRP-secondary anti-rabbit antibody. (D) Hematoxylin, anti-*P.a.* primary antibody. (E) H&E staining of section shown in (A). (F) High magnification of (E).

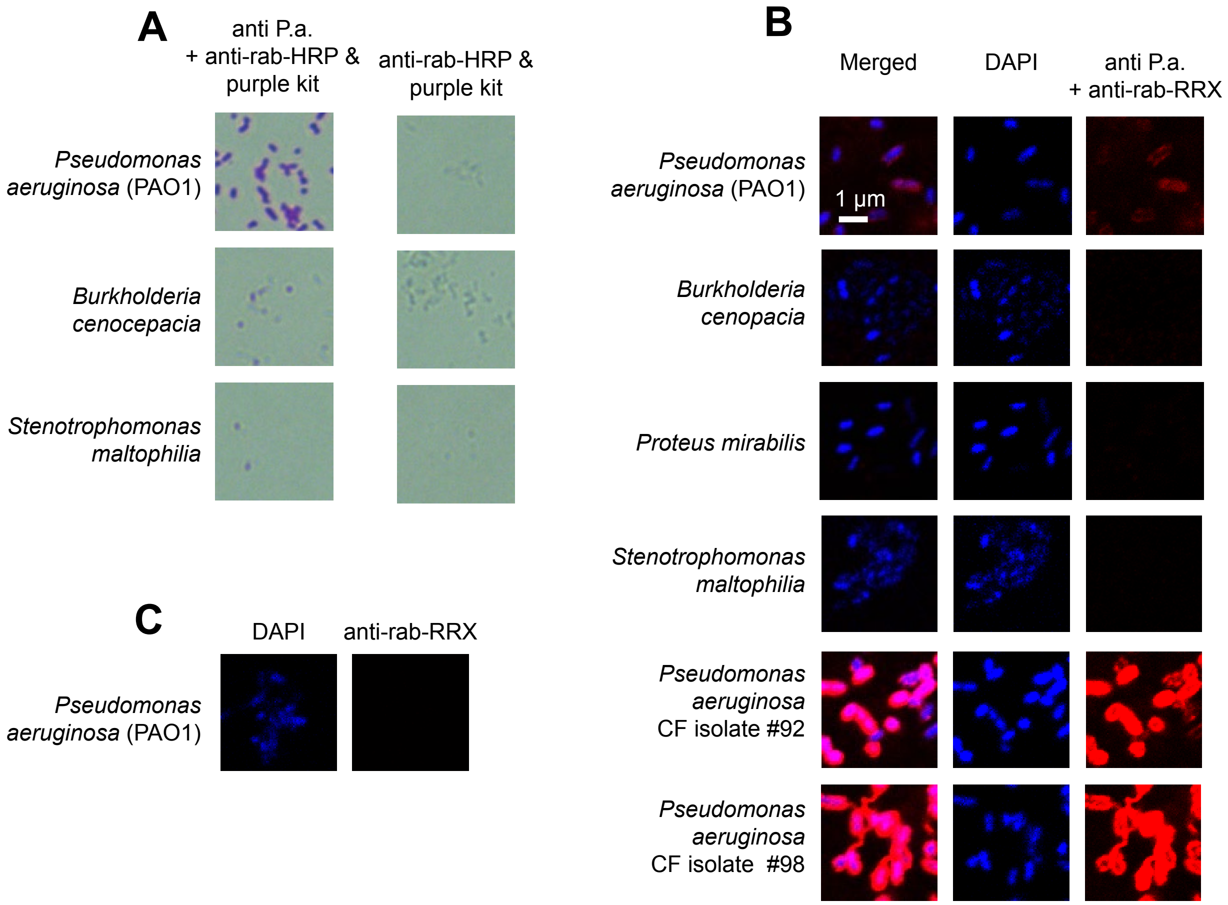

**Supplementary figure E2. Staining controls for antibody specificity using bacterial culture.** Bacterial culture were embedded in paraffin, sliced and processed for IHC using primary and secondary or secondary antibody alone and analysed with a transmission microscope. Alternatively, bacterial suspensions of the indicated strains were fixed and stained using (B) anti-P.a. primary antibody and secondary anti rabbit or (C) the secondary antibody alone and DAPI and analyzed by confocal microscopy.

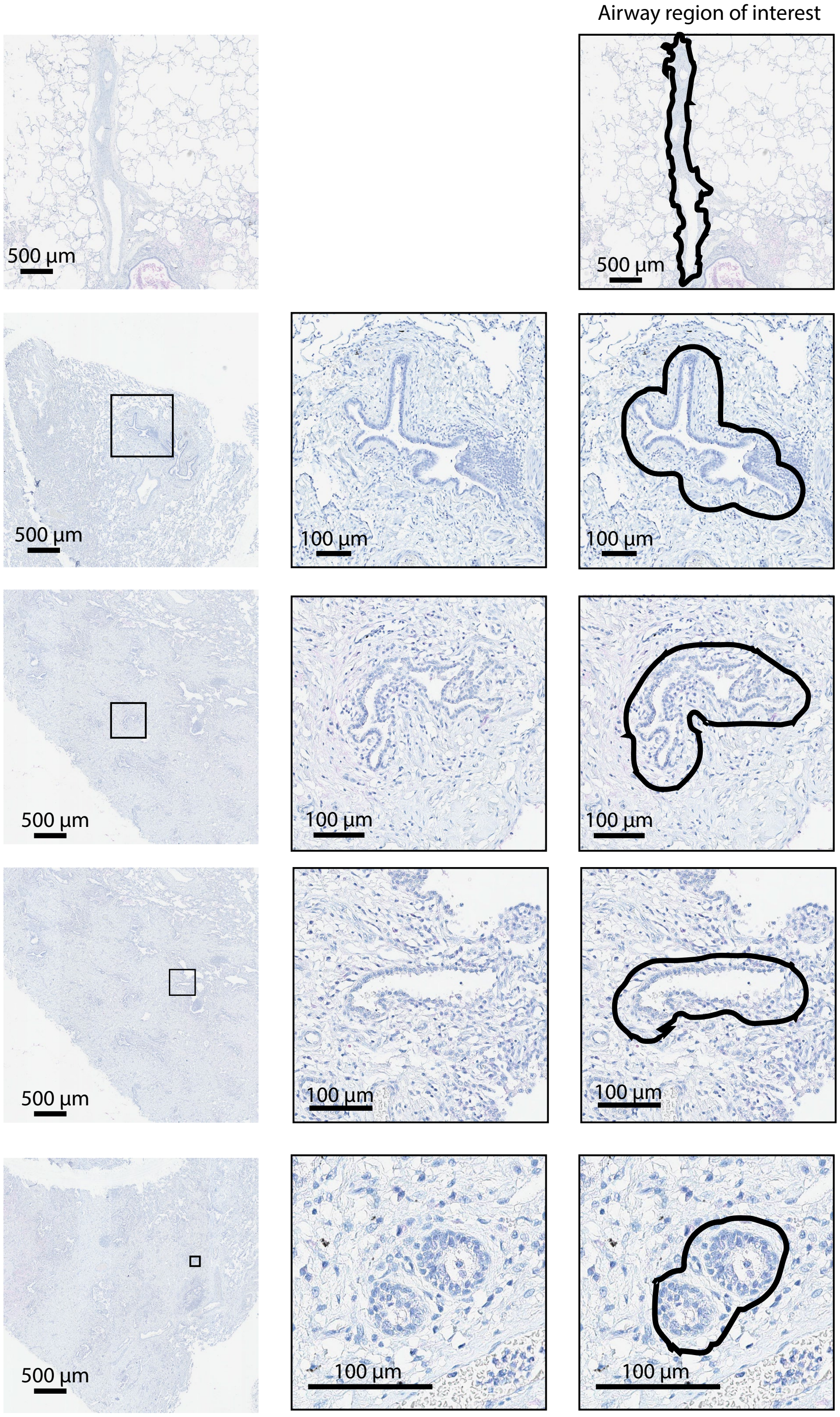

**Supplementary figure E3. Representative microscopy images of different airway sizes analyzed by morphometry and IHC.** Lung tissue thin sections were stained with hematoxylin and anti-*P.a.* antibody for IHC. Left column: low magnification view of tissue sections. Middle column: high magnification view of corresponding marked areas. Right column: region of interest for airway perimeter measurement and detection of intracellular *P.a.* The selected sections represent examples of airway cross-sections whose perimeter range from 38827 to 776 µm.

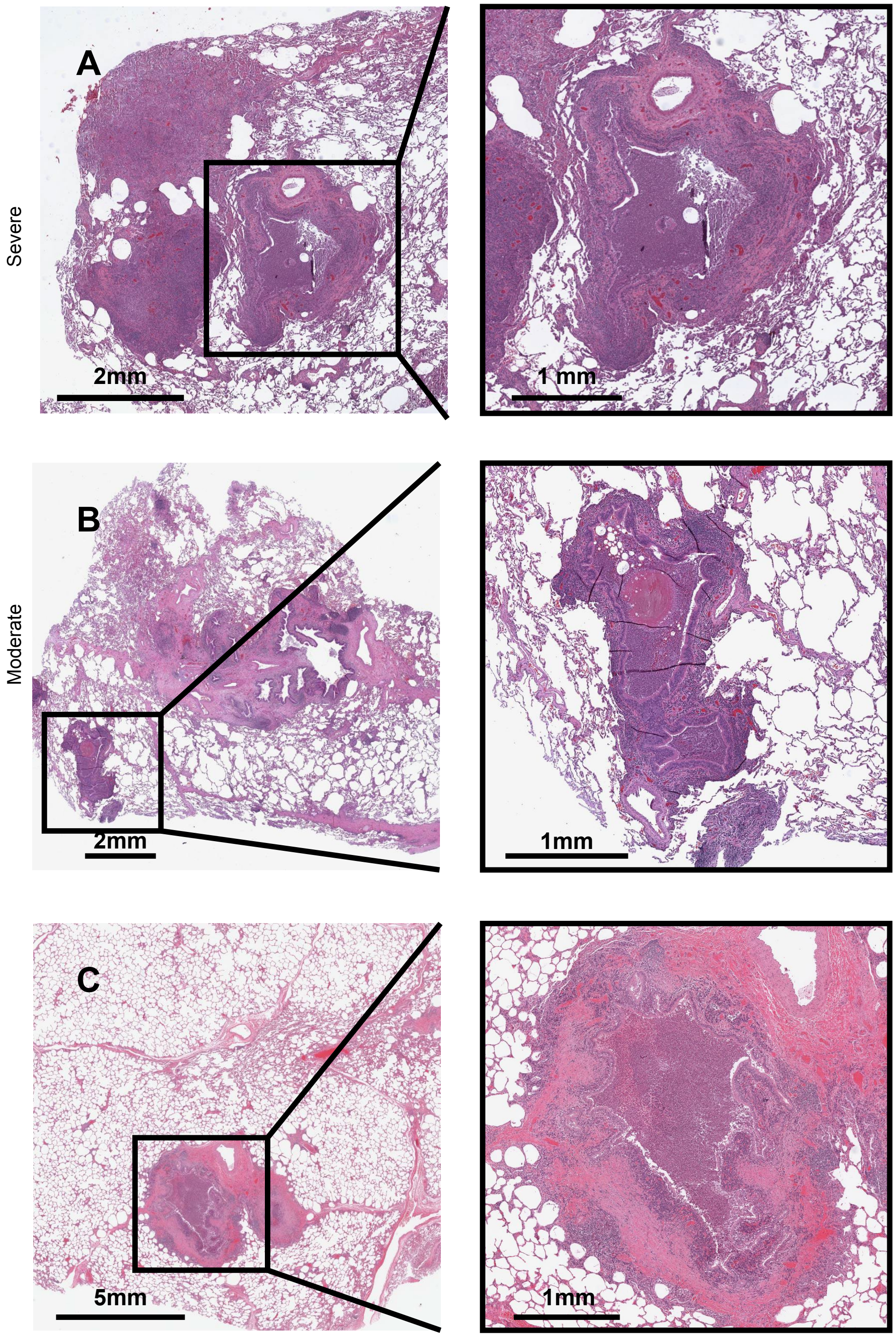

**Supplementary figure E4. Representative histology images of tissue sections used for histopathology scoring of disease severity.** Lung tissue thin sections were stained with H&E and were scored using a semi-quantitative score for inflammation and pathology severity. Left column: low magnification view of tissue sections. Right column: high magnification view of corresponding marked areas. The selected sections represent examples of airway with (A) severe disease (histopathological score of 3) or (B-C) moderate disease (histopathological score of 2).

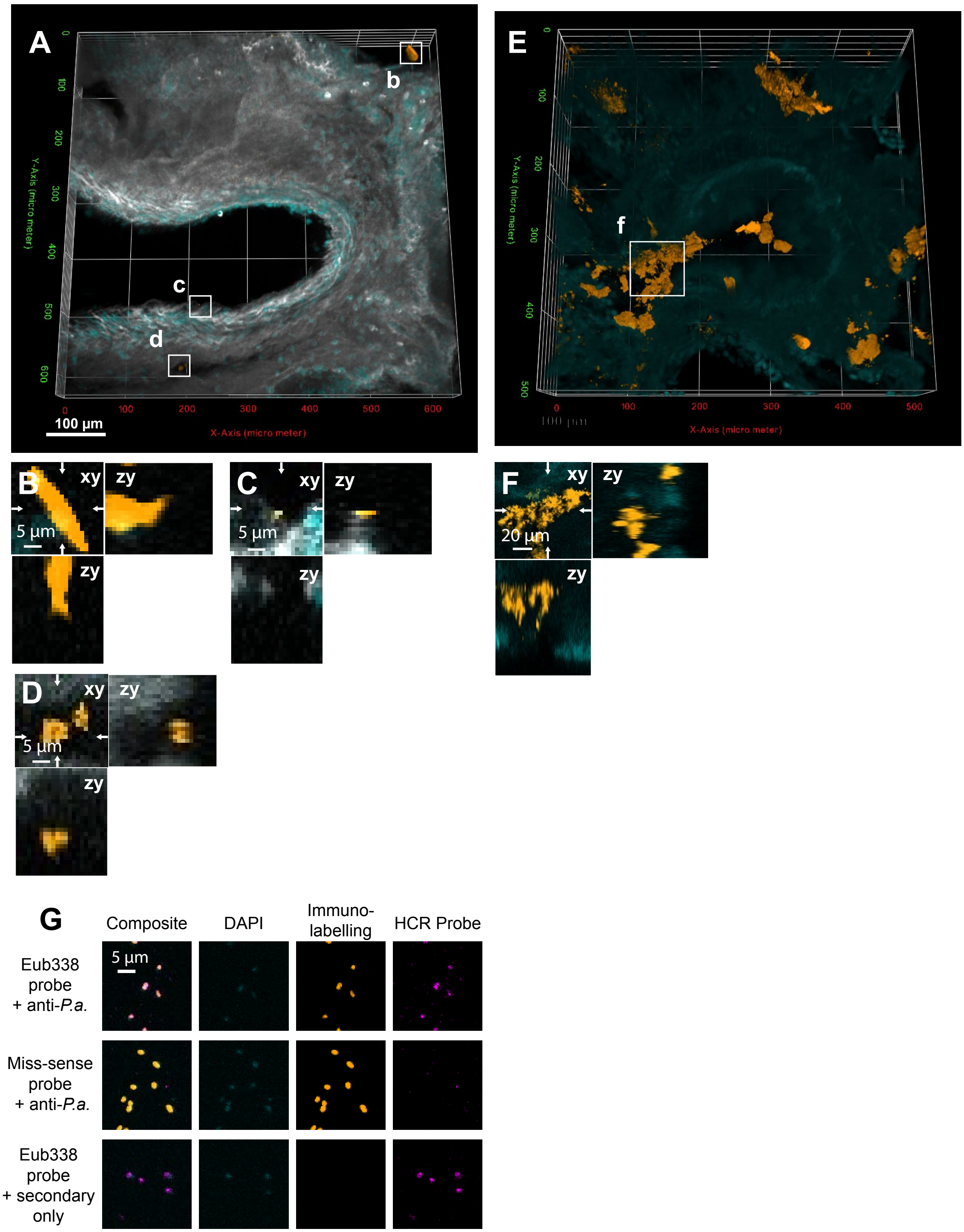

**Supplementary figure E5. Staining controls for MiPACT coupled with immunofluorescence (IF) or hybridization chain reaction (HCR).** (A) Low magnification 3D visualization of explant tissue thick section processed for MiPACT coupled IF control. Staining was done with a secondary anti-rabbit antibody conjugated to Alexa Fluor 488 (orange), Wheat-Germ agglutinin (grey) and DAPI (cyan), but without primary anti-Pa antibody. (B-D) High magnification Orthogonal projections of selected regions (white square area depicting (B) large non-specific aggregates or small aggregates in (C-D) the airway lumen. (E) Low magnification of an alveolar area after staining with a *P.a.*-specific poly-clonal antibody (orange) and DAPI (cyan) and (F) orthogonal projection of the white square area depicting *P.a.* aggregates in the lumen of the alveoli. (G) In vitro HCR and immunofluorescence performed on culture of PA14 strain using the Eub338 probe (magenta) or the non-specific missense probe, the anti-*P.a.* primary antibody with the secondary (orange) or the secondary only and DAPI (cyan).

**Supplementary Movie SM1.** 3D rendering of a CF patient bronchi analyzed by immunofluorescence for *P.aeruginosa.* (orange), DAPI (cyan) and lectin (grey).
